# Optimized Mn²⁺–Phos-tag Gels Reveal Sarcomeric Protein Dephosphorylation upon Myofibril Preparation

**DOI:** 10.64898/2026.08.21.746362

**Authors:** Sunayana Begum Syed, Axel Fenwick, Skylar M. L. Bodt, Rohan Wishard, D. Brian Foster

## Abstract

Precise quantification of myofilament protein phosphorylation is essential for understanding the regulation of cardiac contractility in health and disease. Although Phos-tag SDS-PAGE is widely used to resolve phosphorylated protein isoforms, its reproducibility and quantitative reliability are often limited by variability in the key experimental factors, including gel composition, electrophoretic conditions, protein loading, and sample preparation. Here, we present a standardized Mn²⁺–Phos-tag SDS-PAGE workflow optimized for cardiac myofilament proteins, using myosin regulatory light chain 2 (MLC2) and cardiac troponin I (cTnI) as model targets. We systematically evaluated critical parameters—including Mn²⁺ and Phos-tag concentrations, acrylamide composition, electrophoretic regime, buffer chemistry, protein loading, and EDTA-mediated transfer—to define conditions that maximize phospho-species resolution while preserving quantitative fidelity. We further demonstrate that electrophoresis rate, sample loading, and extraction strategy significantly influence band morphology, signal intensity, and the apparent distribution of phospho-species. As a use case scenario, we compared Trichloroacetic acid (TCA)-extracted mouse left ventricular homogenates with myofibrils prepared using a widely adopted Triton-X-100 tissue-demembranization protocol. Myofibril preparation was associated with profound MLC2 dephosphorylation at the earliest stages of preparation, whereas cTnI exhibited a marked reduction in higher-order, low-stoichiometry phosphoforms. Further evaluation of Myosin-binding protein C (MyBP-C) showed progressive loss of phosphorylation over the course of 24 hours. We submit that TCA-extracted heart standards in combination with Phos-tag gels can provide valuable quality control for the phosphorylation status of myofibril preparations, and that inclusion of a high-affinity PP2A and PP1 phosphatase inhibitor like okadaic acid may benefit future myofibril mechanics studies.

## Introduction

Protein phosphorylation is a fundamental post-translational modification that dynamically regulates enzymatic activity, structural interactions, and signaling across biological systems. In the heart, phosphorylation of myofilament proteins is essential for modulating calcium sensitivity, cross-bridge cycling, and contractile performance during physiological and pathological stress. Cardiac contraction arises from the coordinated interaction of actin-thin and myosin-thick filaments within the sarcomere, where regulatory proteins fine-tune mechanical output to meet changing hemodynamic demands (1). Among the key regulatory proteins, myosin regulatory light chain 2 (MLC2) and cardiac troponin I (cTnI) play central roles in controlling sarcomere activation and relaxation. MLC2 phosphorylation modulates cross-bridge kinetics and thick-filament activation, directly influencing stroke volume and contractile vigor (2–5). CTnI phosphorylation integrates β-adrenergic signaling with thin-filament calcium sensitivity, thereby coordinating lusitropy and systolic force generation (6). Dysregulation of these phosphorylation pathways is recognized as a mechanism contributing to cardiomyopathy, diastolic dysfunction, and heart failure (7–10).

Because MLC2 and cTnI contain multiple phosphorylation sites, they exist as discrete proteoforms, or phospho-species, that correspond to the number of phosphorylated amino acids they bear, each with distinct biophysical consequences. Resolving these proteoforms is, therefore, essential for accurate interpretation of contractile regulation. Phos-tag SDS-PAGE has emerged as a powerful tool for separating phosphorylated proteins based on mobility shifts induced by metal-dependent phosphate affinity (11). In Mn²⁺– or Zn²⁺–Phos-tag gels, phosphorylated residues form stable coordination complexes with the Phos-tag acrylamide ligand, causing precisely graded migration rates proportional to the number of phosphate groups (10, 12–14). This interaction is strictly metal-dependent and is abolished by chelators such as EDTA (15). The resulting coordination chemistry enables resolution of multisite phosphorylation patterns in sarcomeric proteins and has been validated across diverse biochemical studies including cardiac applications (16).

Despite its utility, reproducible Phos-tag analysis of myofilament proteins remains challenging. Band sharpness, phospho-species separation, migration reproducibility, and quantitative interpretation vary markedly between laboratories. These inconsistencies arise from differences in Mn²⁺–Phos-tag concentration, acrylamide percentage, electrophoresis conditions (constant current versus constant voltage), protein loading, reducing agents, gel composition, sample preparation, protein extraction buffers & methods (e.g., RIPA, urea, TCA), chelation efficiency, and transfer conditions. In addition, intrinsic properties of the target protein - including molecular weight, the number and location of phosphorylation sites, phosphorylation stoichiometry, and phosphorylation dependent confirmational changes - can influence Phos-tag binding and electrophoretic mobility. Proteins with closely spaced phosphorylation states—such as MLC2 and cTnI—are particularly sensitive to these parameters; even small deviations can collapse phospho-bands, broaden peaks, distort mobility, or obscure low-abundance species. As a result, cross-study comparisons remain difficult, and subtle but biologically meaningful phosphorylation changes may be overlooked.

Although Phos-tag SDS-PAGE is widely used in cardiovascular research, there is no standardized workflow optimized specifically for myofilament proteins. Existing studies have introduced important improvements—such as neutral-pH Zn²⁺ gels or diagonal electrophoresis (11, 15, 17–19)—but none provide a systematic evaluation of how electrophoresis conditions, gel chemistry, sample preparation, and loading behavior interact to influence the resolution of multi-phosphorylated MLC2 and cTnI species. We address the gap by detailing a Mn²⁺–Phos-tag SDS-PAGE pipeline optimized for resolving multisite phosphorylation of myofilament proteins. Specifically, we compare commonly used Phos-tag SDS-PAGE conditions across cardiac studies, highlighting substantial variability in gel composition, Mn²⁺ concentration, electrophoretic parameters, and sample preparation strategies, as applied to the low molecular weight cardiac myofilament proteins MLC2 and cTnI.

## Materials and Methods

### Animals and Ethics Approval

All animal procedures were approved by the Johns Hopkins University Institutional Animal Care and Use Committee and conformed to National Institutes of Health (NIH) guidelines for the care and use of laboratory animals. Adult C57BL/6J mice (10–16 weeks of age; both sexes) were anesthetized with isoflurane, and ventricles were rapidly excised, rinsed in ice-cold phosphate-buffered saline (PBS), cryopulverized in liquid nitrogen, and stored at −80 °C until further use. For all biochemical analyses, n = 3 biological replicates (R1–R3) per condition were used. Each experiment was independently repeated, and data are presented as the mean of three animals per group. Statistical analyses are described in the corresponding figure legends.

### Reagents and Buffers

All reagents were obtained from Thermo Fisher Scientific or Sigma-Aldrich unless otherwise specified. Phos-tag™ acrylamide was purchased from FUJIFILM Wako Chemicals (cat. no. AAL-107). All buffers were prepared using ultrapure water (>18 MΩ·cm; Milli-Q, Millipore). Buffer pH was adjusted during preparation and verified before use to ensure optimal performance. Buffers and reagents were either freshly prepared or aliquoted for single use to minimize degradation from repeated freeze–thaw cycles and were used within 1–2 months of preparation. Phos-tag™ acrylamide stock solutions were prepared according to the manufacturer’s instructions and stored under the recommended conditions. To preserve Mn²⁺–Phos-tag coordination during electrophoresis, all buffers used for gel preparation and sample handling were free of EDTA, phosphate, and other metal-chelating agents. Chelating reagents were introduced only during post-electrophoretic processing, as described below. A comprehensive list of all reagents, catalog numbers, and final working concentrations—including MnCl_2_ and Phos-tag concentrations used in gel preparation—is provided in **Supplementary Table 1.**

### Tissue Cryopulverization and TCA Precipitation

Frozen ventricular tissue was cryopulverized under liquid nitrogen using a pre-chilled mortar and pestle. Tissue powder was immediately suspended in ice-cold 10% (w/v) trichloroacetic acid (TCA) containing 10 mM dithiothreitol (DTT) and incubated on ice for 30 min with intermittent vortexing to preserve endogenous phosphorylation states. Proteins were pelleted by centrifugation at 10,000 × g for 5 min at 4 °C and subsequently washed three times with ice-cold diethyl ether to remove residual acid, lipids, and reducing agents. Each wash step included brief vortexing followed by centrifugation at 10,000 × g for 5 min at 4 °C. Following the final wash, pellets were briefly air-dried for 5 – 10 min and resolubilized in a denaturing buffer containing 8 M urea, 2% (w/v) SDS, 100 mM NaCl, 100 mM Tris–HCl (pH 8.0), and 10 mM DTT by vortexing briefly, followed by mixing in an Eppendorf ThermoMixer at room temperature until fully dissolved (20, 21).

### Myofibril Isolation and Protein Extraction

Myofibrillar proteins were enriched using a detergent-based permeabilization and differential centrifugation approach adapted from established cardiac muscle preparation methods (22–24). Briefly, ventricular tissue was incubated in ice-cold isolation solution containing 5.55 mM Na_2_ATP, 7.11 mM MgCl_2_, 2 mM EGTA, 108.1 mM KCl, 8.91 mM KOH, and 10 mM imidazole (pH 7.0), supplemented with 2 mM dithiothreitol (DTT), EDTA-free protease inhibitors, and sodium azide, followed by overnight permeabilization in the presence of Triton X-100 at 4 °C. Notably, myofibril isolation buffers contain chelating agents (e.g., EGTA) and require extended handling, which may influence preservation of phosphorylation states. Following permeabilization, samples were washed extensively in isolation solution to remove detergent and glycerol, then homogenized and subjected to low-speed centrifugation (1,500–2,000 × g, 5 min) to pellet the myofibrils. The myofibrillar pellet was repeatedly washed (2–3 times) and subsequently resuspended in relaxing solution containing 5 mM Na_2_ATP, 5.44 mM MgCl_2_, 0.11 mM CaCl_2_, 10 mM EGTA, 100 mM BES, 10 mM creatine phosphate (CrP), and 21.72 mM potassium propionate (pH 7.0). Samples were collected at defined stages of the preparation, including whole lysate, post-wash fractions, and after prolonged incubation (24 h), to assess the effects of sample processing on preservation of phosphorylation states. Prior to Phos-tag SDS-PAGE, myofibrillar proteins were subsequently subjected to methanol–chloroform precipitation to remove buffer constituents. Briefly, methanol, chloroform, and water were sequentially added, vortexed, and centrifuged to induce phase separation. Proteins were recovered from the interphase, washed with methanol, and pelleted by centrifugation, as previously described (25). The resulting protein pellet was then resuspended in 10% SDS for downstream analysis.

### Sample Preparation

Protein concentration was determined using a bicinchoninic acid (BCA) protein assay kit (Thermo Fisher Scientific, Cat. No. 23225) according to the manufacturer’s instructions. Samples were then diluted to the desired concentration in EDTA-free, non-reducing Laemmli sample buffer (62.5 mM Tris-HCl, pH 6.8, 2% SDS, 10% glycerol, and 0.01% bromophenol blue). Dithiothreitol (DTT) was added separately to a final concentration of 5 mM (from a 1 M stock), and samples were incubated at 55 °C for 10 min with mixing at 700 rpm using an Eppendorf ThermoMixer. Following incubation, samples were briefly vortexed and centrifuged to collect condensate before loading.

### Mn²⁺–Phos-tag SDS-PAGE Gel Preparation

Phos-tag SDS-PAGE was performed using 10% polyacrylamide resolving gels containing Phos-tag™ acrylamide and MnCl_2_. All solutions were prepared using EDTA-free reagents to preserve Mn²⁺–Phos-tag coordination. Glass plates (short and spacer plates; Invitrogen) were cleaned with 70% ethanol and assembled according to the manufacturer’s instructions using 1.0 mm spacers. The bottom and sides of the plates were sealed with molten 1% agarose prepared in deionized water to prevent leakage, and the plates were allowed to solidify completely before gel casting. Resolving gels were prepared using 1.5 M Tris–HCl (pH 8.8), acrylamide/bis-acrylamide (37.5:1) at a final concentration of 10% (w/v), and 0.1% SDS. Phos-tag™ acrylamide and MnCl_2_ were included at final concentrations of 50 µM and 300 µM, respectively. Phos-tag acrylamide, MnCl_2_, and SDS were premixed before incorporation into the gel solution to ensure complete metal–ligand complex formation. Polymerization was initiated by the addition of TEMED (0.05–0.1% v/v final) and freshly prepared ammonium persulfate (APS; 0.1% w/v final). Gels were gently mixed to minimize bubble formation and immediately loaded between the glass plates using a 1mL pipette. The gel surface was overlaid with 100% isopropanol to promote a flat gel interface and uniform polymerization. Following polymerization (∼30–45 min), the isopropanol was carefully removed, and the gel surface was rinsed several times with deionized water to remove residual isopropanol before casting the stacking gel. Stacking gels (4%) were prepared using 1 M Tris–HCl (pH 6.8), acrylamide/bis-acrylamide (37.5:1), and 0.1% SDS without inclusion of Phos-tag™ acrylamide or MnCl_2_. Following the addition of TEMED and freshly prepared ammonium persulfate, the stacking gel solution was loaded immediately on top of the polymerized resolving gel using a 1 mL pipette, and a 1.0 mm comb was inserted carefully to avoid bubble formation. Gels were allowed to polymerize at room temperature for at least 1-2 h. After polymerization, wells were rinsed thoroughly with deionized water to remove residual acrylamide. Gels were either used immediately or stored at 4 °C wrapped in paper towels wetted with distilled-deionized water and used within 24 h for optimal performance. Detailed gel compositions and reagent volumes are provided in **Supplementary Table 1**.

### Electrophoresis & Blotting

Protein electrophoresis was performed using standard Laemmli Tris–glycine–SDS running buffer. Samples were prepared in 1× EDTA-free Laemmli sample buffer and supplemented with reducing agent dithiothreitol (DTT) as described above. For mouse cardiac samples, 30 µg of total protein per lane was loaded for MLC2 analysis and 40 µg per lane for cTnI analysis. Samples were loaded using Hamilton syringes to ensure accurate and reproducible delivery into wells. Gels were initially run at 100 V for 10 - 15 min to allow sample entry into the stacking gel. Wells were then gently rinsed with running buffer to remove residual salts and glycerol prior to continuation of electrophoresis. Gels were then resolved by low-voltage electrophoresis at constant voltage (50 V for MLC2, ∼12 h; 50 V for cTnI, ∼14/16 h) at 4 °C. These conditions minimized gel heating and improved the resolution of phospho-species. Following electrophoresis, gels were incubated three times for 10 min in Towbin transfer buffer (25 mM Tris, 192 mM glycine, 20% methanol, pH ∼8.3) containing 30 mM EDTA, with gentle agitation, to chelate Mn²⁺ and dissociate phosphorylated proteins from the Phos-tag matrix. Gels were subsequently washed three additional times for 10 min in EDTA-free transfer buffer to restore conductivity prior to transfer (14). The volume of buffer used for each wash was adjusted according to container size, ensuring complete submersion of the gel and adequate solution exchange during incubation. Proteins were transferred to 0.2 µm nitrocellulose membranes using a wet tank transfer system (Bio-Rad Laboratories, Hercules, CA, USA) at 4 °C at 120 V for 2 h. To minimize heat generation and maintain consistent transfer efficiency, the transfer was performed at 4 °C with the transfer apparatus packed with ice throughout the run.

### Immunodetection & Quantitation

Following protein transfer, membranes were stained with Revert™ 700 Total Protein Stain (LI-COR) for 5–10 min to verify uniform protein transfer and loading. Membranes were rinsed in wash solution containing 6.7% (v/v) glacial acetic acid and 30% (v/v) methanol in water for 5 min, followed by three washes in deionized water (5 min each), and subsequently imaged using a LI-COR imaging system. Membranes were then incubated in Revert™ destaining (reversal) solution containing 0.1 M sodium hydroxide and 30% (v/v) methanol in water for 5 min. Following destaining, membranes were equilibrated in Tris-buffered saline (TBS; 3 × 5 min) and subsequently in TBS containing 0.1% Tween-20 (TBST; 3 × 5 min). Membranes were blocked in 5% (w/v) bovine serum albumin (BSA) prepared in TBST for 1 h at room temperature or overnight at 4 °C. Unless otherwise indicated, all staining, washing, and incubation steps were performed with gentle rocking. Blocked membranes were incubated overnight at 4 °C with primary antibodies against myosin light chain 2 (MLC2; Abcam, ab92721), cardiac troponin I (cTnI; Sigma-Aldrich, MAB1691), and myosin-binding protein C (cMyBP-C; ProSci, 56-426), each diluted 1:1000 in blocking buffer. Following primary antibody incubation, membranes were washed in TBST (3 × 5 min) and incubated with fluorescent IRDye® 680RD and/or 800CW secondary antibodies (LI-COR; 1:5000 dilution) for 1 h at room temperature. Membranes were subsequently washed again in TBST (3 × 5 min), followed by a final wash in TBS for 5 min, and then transferred to fresh TBS prior to signal detection using the LI-COR Odyssey imaging system.

### Image Analysis

Densitometric analysis of Phos-tag immunoblots was performed using LI-COR Image Studio software. Immunoblots were imaged using a LI-COR Odyssey F imaging system (LI-COR Biosciences) at 169 µm resolution. Images were acquired using the appropriate fluorescence channel(s) under nonsaturating conditions and with consistent acquisition settings across comparable samples to maintain signal detection within the linear range. This was particularly important when high- and low-abundance phospho-species were present within the same lane, as saturation of a predominant phospho-species could lead to underestimation of its intensity and bias the calculated phosphorylation-state distribution. If saturation was detected, the blot was reimaged at a lower acquisition intensity before quantitative analysis.

Phos-tag bands were quantified directly from the original acquired images. Individual phospho-species were defined using rectangular regions of interest (ROIs), with consistent ROI dimensions maintained for comparable bands within an experiment. ROIs were duplicated and repositioned as necessary to preserve consistent dimensions. Local background correction was applied using the Image Studio background-subtraction function, and the resulting background-corrected signal values were used for quantitative analysis. Brightness and contrast adjustments, when required for visualization, were applied uniformly to the entire image and were not applied to images used for densitometric quantification. For each sample, the background-corrected intensities of all resolved phospho-species within a lane were summed to obtain the total protein signal. The relative abundance of each phosphorylation state was calculated as:

Phospho-species (%) = [intensity of individual phospho-species / sum of intensities of all phospho-species within the same lane] × 100.

### Preparation of Negative Control Phospho-Standards (Alkaline Phosphatase Treatment)

Protein lysates were incubated with calf intestinal alkaline phosphatase (CIP; Thermo Fisher Scientific, Cat. No. 18009027) at approximately 1 µg protein per unit enzyme to achieve complete dephosphorylation. Reactions were performed in a total volume of ∼25 µL, containing CIP, 10× reaction buffer, and protein lysate, with deionized water added as required. Enzyme and buffer conditions were empirically optimized to ensure maximal phosphatase activity. Samples were incubated for 16–18 h at 37 °C with gentle agitation (∼700 rpm), and reactions were subsequently terminated by heating at 65 °C for 10 min according to the manufacturer’s instructions. Samples were then stored at −80 °C until further use. Prior to electrophoresis, samples were mixed with Laemmli sample buffer supplemented with DTT and heated at 95 °C for 5 min. Equal protein loading was maintained across all samples. This treatment resulted in efficient conversion of phosphorylated species into the corresponding non-phosphorylated (P0) forms and was used as a negative phospho-control for Phos-tag SDS-PAGE analyses.

### Phos-tag Quantification and Statistical Analysis

Phosphorylation states of proteins resolved by Mn²⁺–Phos-tag SDS-PAGE were quantified using background-corrected signal intensities obtained with LI-COR Image Studio software, as described above. For each sample, the intensity of individual phospho-species (e.g., P0–P3 for MLC2; P0–P5 for cTnI; and P0–P2 for cMyBP-C) was expressed as a percentage of the total signal from all resolved phospho-species within the corresponding lane. Sample-wise phospho-state distributions were verified to total approximately 100% following normalization. All downstream data processing, statistical analyses, and visualization were performed in R using the tidyverse package. Data are presented as mean ± SEM, with individual biological replicates shown where applicable. Statistical comparisons were performed independently for each phosphorylation state using paired or unpaired two-tailed t-tests, as appropriate to the experimental design. Where multiple comparisons were performed, p-values were adjusted using the Benjamini–Hochberg method, with adjusted p < 0.05 considered statistically significant. Phosphorylation-state distributions were visualized using stacked bar plots, while individual phosphorylation states were displayed as mean ± SEM with individual biological replicates overlaid. A comparison of representative Mn²⁺–Phos-tag SDS-PAGE protocols and a troubleshooting guide for the optimized protocol are provided in **Tables 1 and 2**, respectively.

## Results

### Overview of the optimized Mn^2+^–Phos-tag workflow

Our optimized Mn^2+^–Phos-tag workflow is summarized in **Figure 1A**. Briefly, ventricular tissue was flash-frozen in liquid nitrogen and cryopulverized followed by protein extraction using ice-cold trichloroacetic acid (TCA) containing 10 mM dithiothreitol (DTT) to preserve endogenous phosphorylation states prior to electrophoresis and immunoblotting. Myosin light chain 2 (MLC2) and cardiac troponin I (cTnI) were selected as benchmark substrates due to their well-characterized, multisite phosphorylation patterns and established central roles in cardiac contractile regulation (**Figure 1B-G**). These proteins provide robust and physiologically relevant models for evaluating phospho-state resolution using Phos-tag SDS-PAGE. Although MLC2 and cTnI have predicted molecular weights of ∼19 kDa and ∼24 kDa, respectively, band positions in Mn^2+^–Phos-tag gels reflect phosphorylation-dependent mobility shifts rather than true molecular weight. Accordingly, the unphosphorylated MLC2 (P0) species migrated between ∼15–20 kDa, while mono-, di-, and tri-phosphorylated bands (designated P1, P2, and P3) respectively appeared at ∼25 kDa. Similarly, cTnI P0 migrated near ∼25 kDa, whereas highly phosphorylated species (e.g., P4 & P5) exhibited markedly reduced mobility and appeared at apparent molecular weights approaching ∼75 kDa. These shifts arise from the transient binding of phosphate groups to the Phos-tag–Mn^2+^ complex, which retards electrophoretic migration in proportion to phosphorylation state. To verify that the bands above P0 are indeed phosphoforms, MLC2 was treated with alkaline phosphatase, which removed bands P1-P3 (**Figure 1B–D**). Conversely, phosphorylation of cTnI was enhanced via β-adrenergic stimulation of mouse hearts with isoproterenol (ISO) prior to protein extraction (**Figure 1E–G**). These complementary perturbations enabled assessment of both loss and gain of phosphorylation.

**Figure 1.**
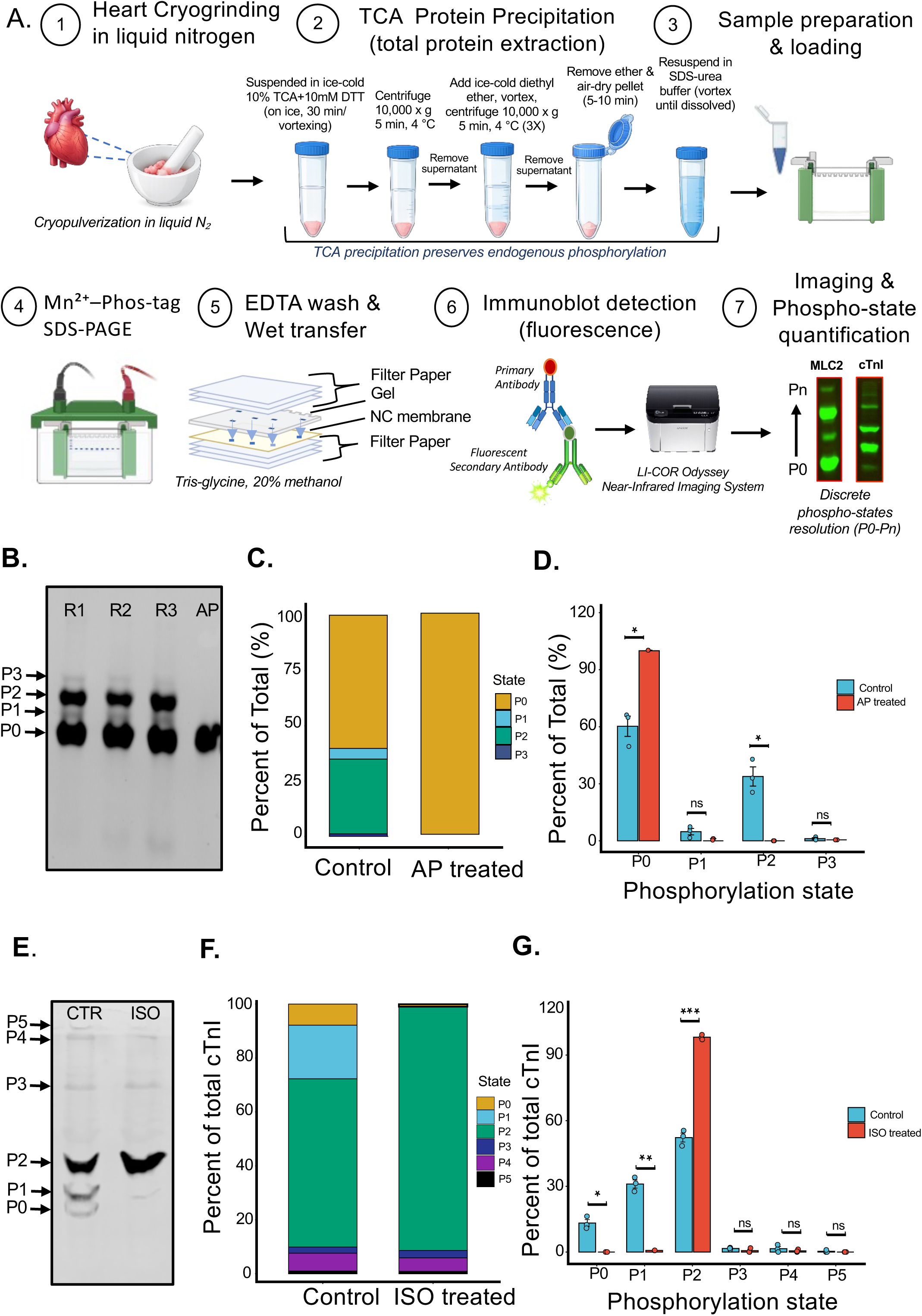
Mn^2+^–Phos-tag workflow and validation using cardiac phosphoproteins. (A) Schematic overview of the optimized Mn^2+^–Phos-tag SDS-PAGE workflow. Ventricular tissue is cryopulverized under liquid nitrogen, and proteins are extracted using ice-cold trichloroacetic acid (TCA) containing 10 mM dithiothreitol (DTT) to preserve endogenous phosphorylation. Samples are resolved on Mn^2+^–Phos-tag gels, subjected to EDTA prewashes before transfer, and analyzed by immunoblotting and quantitative imaging. (B) Representative Phos-tag immunoblot of MLC2 showing phosphorylation-dependent separation into discrete phospho-species (P0–P3). Alkaline phosphatase (AP) treatment collapses higher-order phospho-species into the non-phosphorylated P0 band, confirming phosphorylation-dependent mobility. R1–R3 denote biological replicates. (C) Average phospho-state distribution of MLC2 comparing control and AP-treated samples, expressed as percentage of total signal per lane. AP treatment results in near-complete redistribution to the P0 state. (D) Quantification of individual MLC2 phospho-states (P0–P3). AP treatment significantly increases P0 and reduces higher-order phospho-species. Data are presented as mean ± SEM (n = 3 biological replicates); *p < 0.05; ns, not significant. (E) Representative Phos-tag immunoblot of cardiac troponin I (cTnI) under control (CTR) and isoproterenol (ISO) treatment. ISO stimulation induces a shift toward higher-order phospho-species (P2–P5). (F) Average phospho-state composition of cTnI under control and ISO-treated conditions. (G) Quantitative distribution of cTnI phospho-states (P0–P5). ISO treatment significantly decreases lower phosphorylation states (P0, P1) and increases higher phosphorylation states, particularly P2. Data are presented as mean ± SEM (n = 3 biological replicates); *p < 0.05, **p < 0.01, ***p < 0.001; ns, not significant. Apparent molecular weights reflect phosphorylation-dependent mobility shifts in Phos-tag gels rather than true protein size.

### Optimization of Phos-tag gel chemistry for maximum resolution

To identify gel conditions that provide maximal resolution of MLC2 phosphorylation states, we systematically varied MnCl_2_ concentration, Phos-tag reagent concentration, and acrylamide percentage—three key parameters that collectively influence metal–phosphate coordination and electrophoretic mobility. Although these factors are individually known to influence Phos-tag performance, their combined effects on multi-phosphorylated myofilament proteins have not been systematically evaluated. Mn^2+^ concentration strongly influenced both phosphate-dependent and phosphate-independent mobility of MLC2 (**Figure 2A–D**). High Mn^2+^ (1mM) greatly slowed migration of all bands, including the phosphate-free proteoform (**Figure 2B**), compared to the gels run in 600 µM Mn^2+^ (**Figure 2C**) and 300 µM Mn^2+^ (**Figure 2D**). Consistent with this observation, the relative mobility (Rf) of the unphosphorylated band increased as Mn^2+^ concentration decreased, enabling improved entry into the resolving gel. Greater band mobility in 300 µM Mn^2+^ permitted substantially better separation of the phosphoforms, while reducing smearing (**Figure 2E-F**). Using 300 µM Mn^2+^ gels as a foundation, the greatest impact on phosphoform resolution came from raising the concentration of Phos-tag reagent for MLC2 (**compare Figure 2D with 2H**). Raising the concentration from 30 µM (**Figure 2D**) to 50 µM (**Figure 2H**) permitted full resolution of phosphoforms (P1-P3). Further raising the concentration to 75 µM (**Figure 2I**) only increased separation marginally. Finally, we varied the acrylamide percentage from 10 to 12%. Though 12% gels increase band sharpness, the tradeoff was slower band migration and, consequently, more modest phosphoform separation. We opted to use 10% for further analysis, though 11% (**Figure 2J**) and 12% (**Figure 2K, L**) gels remain a viable option.

**Figure 2.**
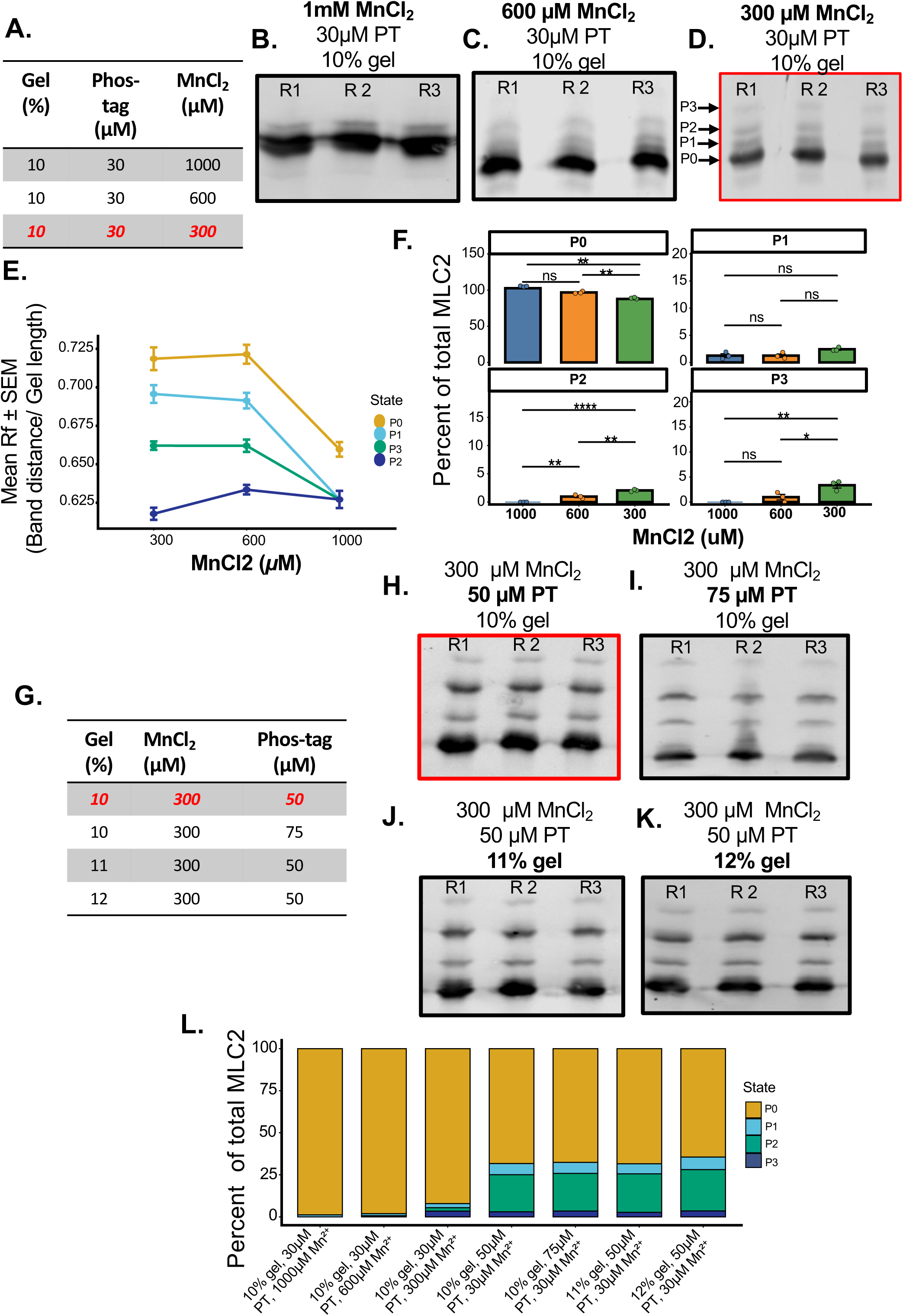
Optimization of Mn^2+^–Phos-tag gel chemistry defines conditions for high-resolution separation of MLC2 phosphorylation states. (A) Summary of the MnCl_2_ concentrations evaluated for Mn^2+^–Phos-tag SDS-PAGE optimization using a fixed gel composition (10% acrylamide, 30 µM Phos-tag). (B–D) Representative Mn^2+^–Phos-tag immunoblots of MLC2 phosphorylation states (P0–P3) resolved using 1 mM (B), 600 µM (C), and 300 µM (D) MnCl_2_. High Mn^2+^ concentrations produced excessive migration retardation and compression of phosphoforms, whereas 300 µM MnCl_2_ enabled improved separation of individual phosphorylation states. R1–R3 denote biological replicates. (E) Quantification of the relative electrophoretic mobility (Rf) of individual MLC2 phosphorylation states as a function of MnCl_2_ concentration. Data are presented as mean ± SEM. Increasing Mn^2+^ concentration reduced protein mobility and compressed phosphoform separation, particularly for the higher phosphorylation states. (F) Quantification of individual MLC2 phosphorylation states (P0–P3) across MnCl_2_ concentrations. While P0 and P1 remained unchanged, the abundance of the P2 and P3 phosphoforms increased significantly at 300 µM MnCl_2_, reflecting improved resolution of higher-order phosphorylation states. Data are presented as mean ± SEM (n = 3 biological replicates); *p* < 0.05; ns, not significant. (G) Summary of Phos-tag concentrations and acrylamide percentages evaluated using the optimized MnCl_2_ concentration (300 µM). (H–K) Representative Mn^2+^–Phos-tag immunoblots illustrating the effects of increasing the Phos-tag concentration (50–75 µM) and acrylamide concentration (10–12%) on MLC2 phosphoform resolution. Increasing the Phos-tag concentration to 50 µM in a 10% gel further improved phosphoform separation, whereas increasing the Phos-tag concentration to 75 µM or the acrylamide concentration to 11–12% did not further improve resolution and resulted in modest band compression. (L) Distribution of MLC2 phosphorylation states (P0–P3) under each gel condition, expressed as the percentage of total signal per lane. Although the overall phosphorylation-state distribution remained largely conserved, gel chemistry influenced the apparent distribution of individual phosphoforms by altering their electrophoretic resolution. Collectively, these data identify 300 µM MnCl_2_, 50 µM Phos-tag, and a 10% acrylamide gel as the optimal conditions for resolving MLC2 phosphorylation states.

### Impact of Sample Preparation, Handling, & Loading

Prior work has indicated that MLC2 undergoes rapid dephosphorylation upon cardiac tissue homogenization, even when a strong denaturant such as SDS is included in the homogenization buffer. The same study found that only ice-cold TCA precipitation preserved maximum phosphorylation (26, 27). We wanted to evaluate whether rapid methanol/water/chloroform precipitation using the method of Wessel & Flugge (25), could preserve as effective as TCA precipitation, as the extraction method likewise serves as a convenient and effective protein clean-up step prior to electrophoresis. TCA precipitation provided more consistent protein recovery and improved resolution of MLC2 phospho-species compared with methanol-based precipitation (**Figure S1A**). However, resolubilizing TCA-extracted denatured protein can be challenging. We therefore tested solubilization strategies for TCA-precipitated samples; an SDS–urea–based buffer (8 M urea, 2% SDS, 100 mM NaCl, 100 mM Tris–HCl, pH 8.0) enabled complete protein solubilization and optimal band resolution, whereas alternative buffers resulted in incomplete solubilization or distorted migration (**Figure S1B, C**). We also evaluated the impact of buffer composition on alkaline phosphatase treatment efficiency and apparent Phos-tag mobility. The observed differences were consistent with variations in compatibility with Mn^2+^–Phos-tag coordination (**Figure S1D**).

Mindful that proteins undergo heat-catalyzed phosphate hydrolysis and/or b-elimination, we further assessed the impact of SDS sample heating temperature and incubation time on MLC2 phosphostatus (**Figure 3A**). Quantitative analyses revealed only modest temperature-dependent effects, with small reciprocal changes in P0 and P1, stable P2 levels, and consistently low P3 abundance (**Figure 3B**). High-temperature denaturation (95 °C) resulted in protein degradation, whereas moderate conditions (55 °C) produced optimal band resolution. In contrast, lower temperature denaturation (37 °C) resulted in reduced band resolution (data not shown).

**Figure 3.**
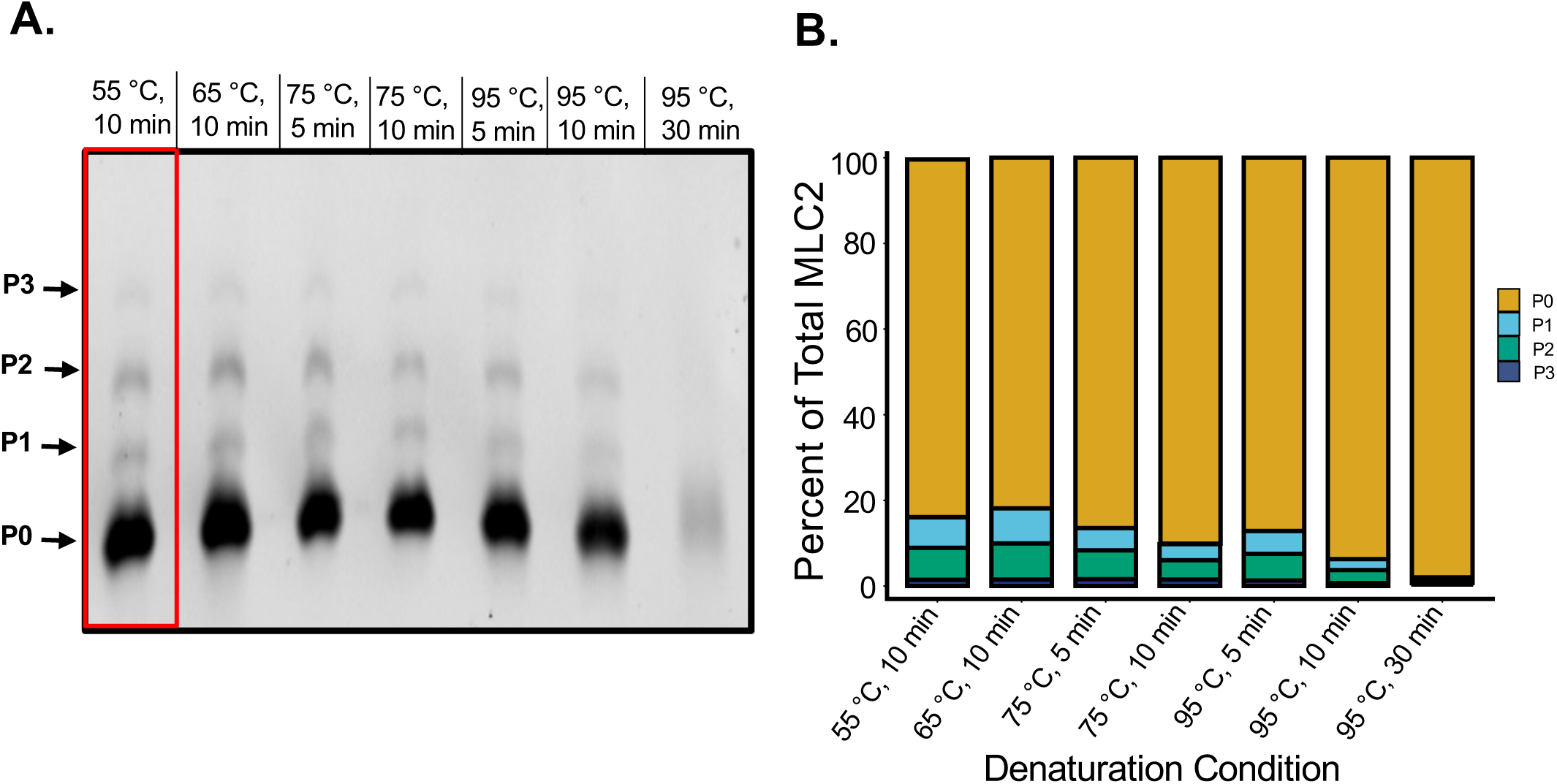
Sample denaturation conditions influence the apparent phosphorylation-state distribution of MLC2 in Mn^2+^–Phos-tag SDS-PAGE. (A) Representative Mn^2+^–Phos-tag immunoblot of ventricular myosin light chain 2 (MLC2) following sample denaturation under the indicated temperature and incubation conditions (55°C for 10 min, 65°C for 10 min, 75°C for 5 or 10 min, and 95°C for 5, 10, or 30 min). Phosphorylation states (P0–P3) are indicated on the left. Increasing the denaturation temperature and incubation time progressively reduced the detection of higher-order phosphoforms, with the greatest loss observed after incubation at 95 °C for 30 min. The 55 °C for 10 min denaturation condition (red box) was selected for all subsequent experiments because it best preserved the apparent MLC2 phosphorylation-state distribution. (B) Distribution of MLC2 phosphorylation states (P0–P3), expressed as the percentage of the total MLC2 signal per lane across the indicated denaturation conditions. Higher denaturation temperatures shifted the apparent phosphorylation-state distribution toward the unphosphorylated (P0) state, indicating reduced preservation of phosphorylated MLC2 under harsh denaturation conditions.

We next examined the influence of sample loading on Phos-tag band resolution and phosphorylation-state quantification by analyzing MLC2 and cTnI across increasing protein loads and quantifying phospho-state distributions (**Figure 4A–F**). For MLC2, increasing the loading from 5 to 20 µg preserved clear resolution of the P0–P3 species (**Figure 4A**), and quantification demonstrated that the overall phospho-state distribution remained largely stable across loading conditions (**Figure 4B**). State-wise analysis revealed modest, load-dependent shifts within individual phospho-species, including a slight decrease in P0 and corresponding increase in P2 at higher loading (**Figure 4C**), consistent with the observed band patterns. In contrast, cTnI exhibited a pronounced loading-dependent increase in apparent phosphorylation-state complexity. Increasing protein loading from 20 to 80 µg resulted in stronger signal intensity and enhanced detection of higher-order phospho-species (**Figure 4D**). Despite these changes, stacked distributions indicated that the overall fractional distribution across P0–P5 remained broadly consistent across loading conditions (**Figure 4E**). State-wise analyses (**Figure 4F**) revealed load-dependent differences that primarily reflect changes in band resolution and detectability, rather than true alterations in phosphorylation stoichiometry. Accordingly, these effects did not substantially alter the overall phosphorylation balance.

**Figure 4.**
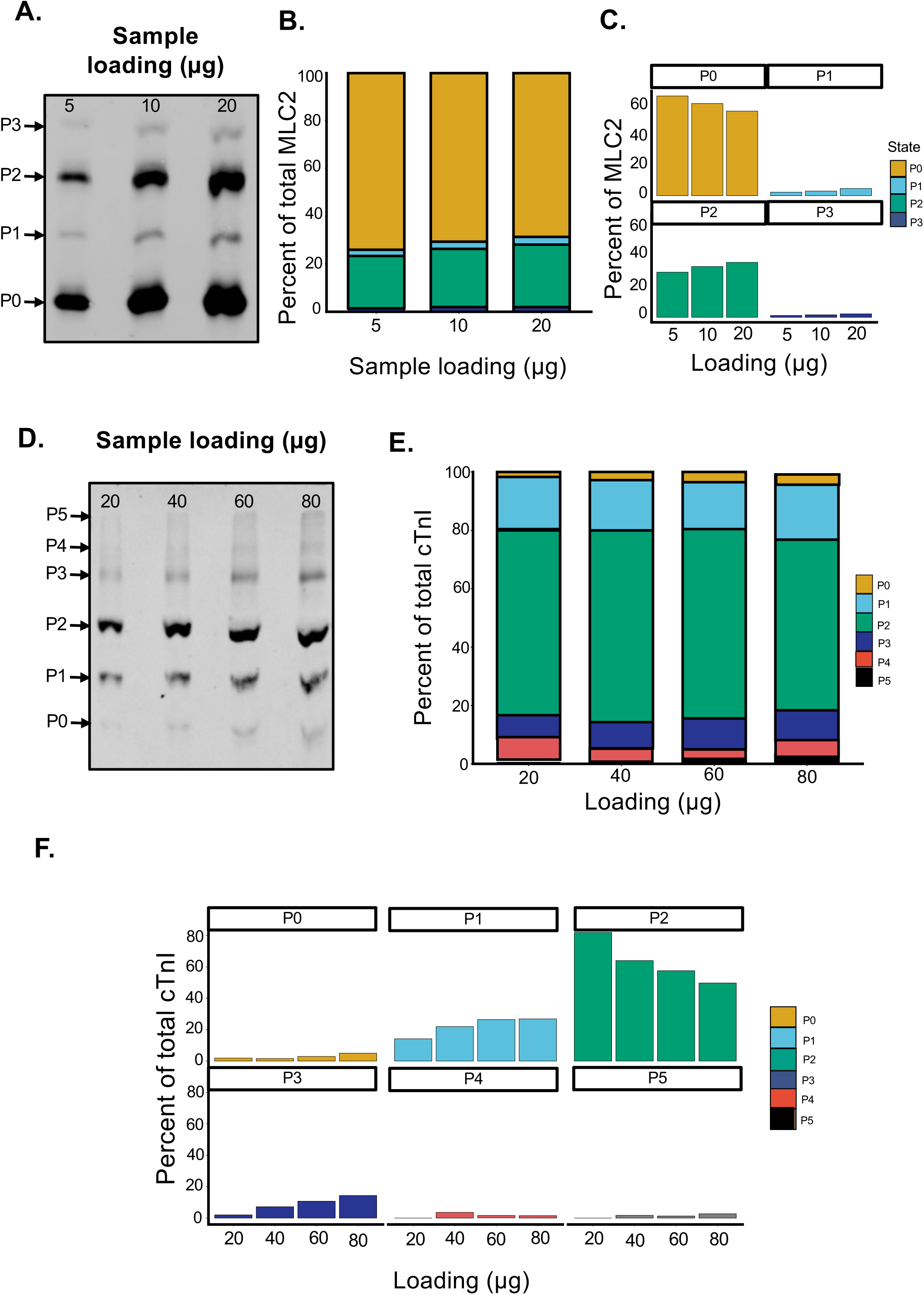
Protein loading influences phosphoform detection and quantification of MLC2 and cTnI in Mn^2+^–Phos-tag SDS-PAGE. **(A)** Representative Mn^2+^–Phos-tag immunoblot of ventricular myosin light chain 2 (MLC2) resolved at increasing protein loads (5, 10, and 20 µg). Phosphorylation states (P0–P3) are indicated. **(B)** Distribution of MLC2 phosphorylation states (P0–P3), expressed as the percentage of the total MLC2 signal per lane across the indicated protein loading conditions. **(C)** Quantification of individual MLC2 phosphorylation states (P0–P3) across protein loading conditions. Increasing protein loading resulted in modest changes in the relative abundance of individual phosphoforms, primarily involving P0 and P2, while the overall phosphorylation-state distribution remained largely preserved. **(D)** Representative Mn^2+^–Phos-tag immunoblot of cardiac troponin I (cTnI) resolved at increasing protein loads (20, 40, 60, and 80 µg), showing phosphorylation states (P0–P5). **(E)** Distribution of cTnI phosphorylation states (P0–P5), expressed as the percentage of the total cTnI signal per lane across the indicated protein loading conditions. **(F)** Quantification of individual cTnI phosphorylation states (P0–P5) across protein loading conditions. Increasing protein loading enhanced detection of lower-abundance phosphoforms, particularly the higher-order P3–P5 species, while the dominant P2 phosphoform remained relatively stable across loading conditions. Collectively, these data demonstrate that protein loading influences phosphoform detection and the apparent phosphorylation-state distribution in Mn^2+^–Phos-tag SDS-PAGE, emphasizing the importance of standardized protein loading for accurate quantitative comparisons.

Consistent with these findings, complementary analyses (**Figure S2**) further demonstrate that increasing sample loading enhances detection sensitivity and apparent phospho-state complexity without fundamentally altering the overall phosphorylation-state distribution. Together, these results indicate that sample loading influences band morphology and apparent state complexity in Phos-tag gels, while largely preserving the overall phosphorylation-state balance, particularly for MLC2.

### Electrophoresis parameters impact phospho-species resolution

Electrophoresis conditions influenced the apparent resolution and quantification of MLC2 phosphorylation states on Phos-tag gels. Band migration and phospho-state distribution were compared under three commonly used running conditions: constant current (25 mA and 50 mA) and constant low voltage (50 V) (**Figure 5A–C**). Constant current runs minimized band dispersion and offered somewhat greater resolution than at a constant 50V, though separation could be increased through extended run times, allowing the lower molecular weight marker (∼10 kDa) to migrate off the gel to improve separation of protein phosphoforms **(Figure 5A-D)**. Notably, however, low-voltage electrophoresis (50 V for 12 h/ overnight at 4 °C) appeared to maximize the P2 to P0 ratio relative to 25 mA and 50 mA runs (**Figure 5E)** and was therefore used for subsequent experiments.

**Figure 5.**
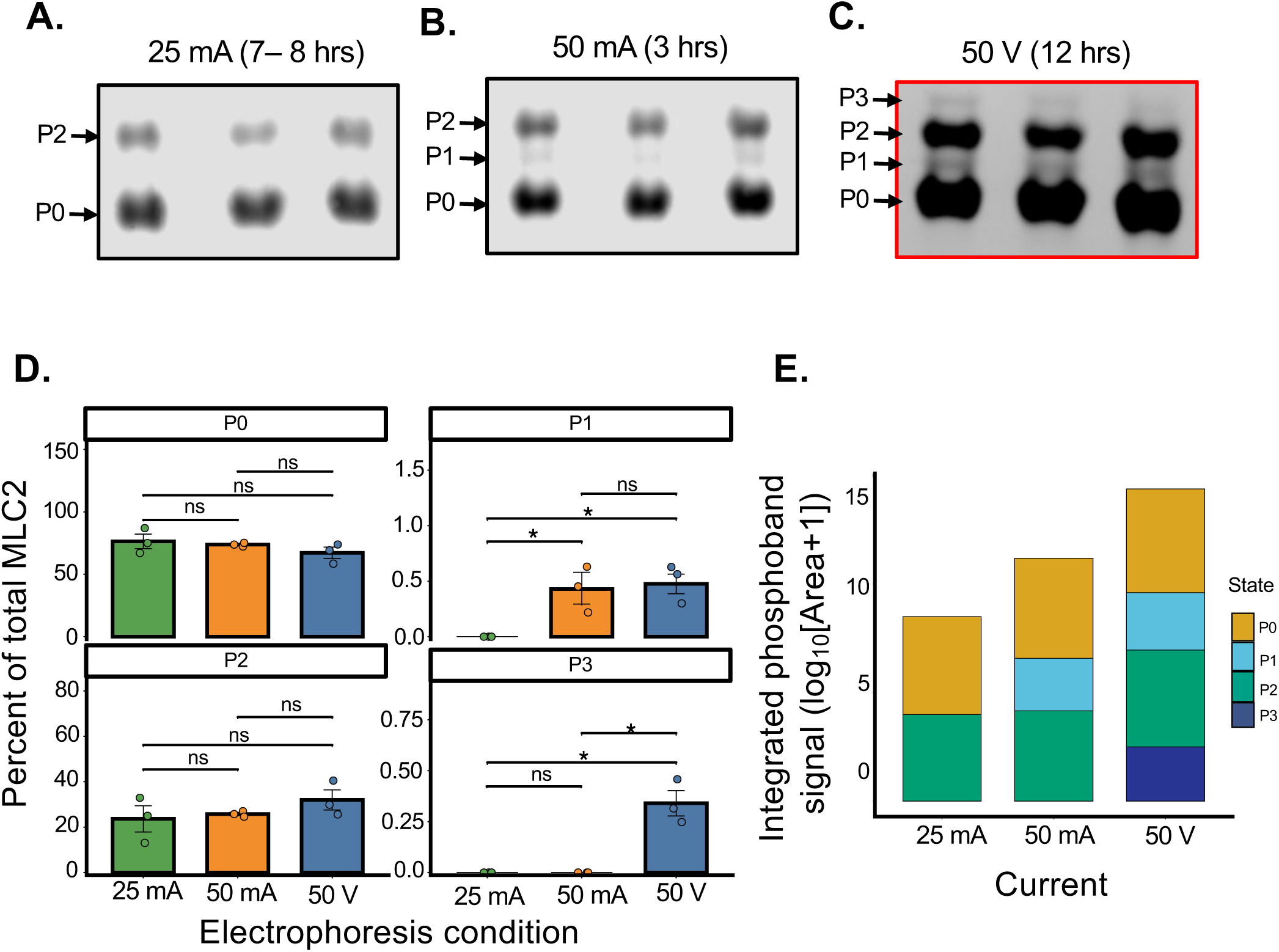
Electrophoresis conditions influence the apparent resolution and quantification of MLC2 phosphorylation states in Mn^2+^–Phos-tag SDS-PAGE. **(A–C)** Representative Mn^2+^–Phos-tag immunoblots of MLC2 showing separation of phosphorylation states (P0–P3) under different electrophoresis conditions: constant current at 25 mA for 7–8 h **(A)**, 50 mA for 3 h **(B)**, and constant voltage at 50 V for 12 h. **(C)**. Prolonged low-voltage electrophoresis increased phosphoform separation compared with constant-current conditions. **(D)** Quantification of individual MLC2 phosphorylation states (P0–P3), expressed as the percentage of the total MLC2 signal (mean ± SEM). Although prolonged low-voltage electrophoresis improved phosphoform separation, the relative abundance of individual phosphorylation states did not differ significantly among the electrophoresis conditions. **(E)** Integrated phosphoband signal of resolved MLC2 phosphoforms (P0–P3), expressed as log₁₀(area + 1), under each electrophoresis condition. Collectively, these results demonstrate that electrophoresis conditions substantially influence the resolution of MLC2 phosphoforms on Mn^2+^–Phos-tag gels. The 50 V for 12 h condition provided the greatest phosphoform separation and was therefore used for all subsequent experiments.

### Evaluation of myofilament protein phosphostatus from a standard myofibril preparation

Given the susceptibility of MLC2 to dephosphorylation under most protein extraction conditions, we evaluated its phosphostatus over the course of the myofibril isolation using a standard method in combination with our refined Phos-tag SDS-PAGE protocol (**Figure 6A-C**). We observed striking differences in phospho-MLC2 status between the myofibrils and TCA-extracted LV homogenates. Notably, even at the earliest stage of myofibril isolation, MLC2 underwent an abrupt and substantial loss of the P2 (di-phosphorylated) band with an initial concomitant increase in the intensity of the P1 (mono-phosphorylated) band, which increased further when checked after the initial Triton X-100 washing phase. After 24 hrs in Triton-skinning solution, marginal levels of the P2 band were detected, and P1 also declined, leaving MLC2 about 95% unphosphorylated. This contrasts with TCA-extracted LV, which was 49% unphosphorylated, 49% di-phosphorylated, and about 2% mono/tri-phosphorylated (**Figure 6C**). CTnI, in contrast, exhibited a loss of phosphorylation, albeit with a distinct pattern. While the P2 state remained relatively stable across fractions, higher-order phosphoforms (P3–P5) were present in TCA samples but markedly reduced in myofibril preparations (**Figure 6D–F**). Consistent with this, quantification of higher-order phosphorylation revealed a significant decrease in P3–P5 across myofibril fractions (**Figure S3A**). Finally, although our Phos-tag protocol was optimized for 20-30 kDa proteins, the distinct character of MLC2 and cTnI phosphorylation loss prompted us to further examine the phosphostatus of cardiac Myosin Binding Protein C (cMyBP-C3; ∼140 kDa) using low-percentage (6%) gels with extended run times (at 50 V) to improve separation of higher-order phosphoforms. CMyBP-C exhibited reduced abundance of higher-order phosphorylation states, particularly P2, in myofibril samples compared to TCA-extracted controls (**Figure S3B–D**).

**Figure 6.**
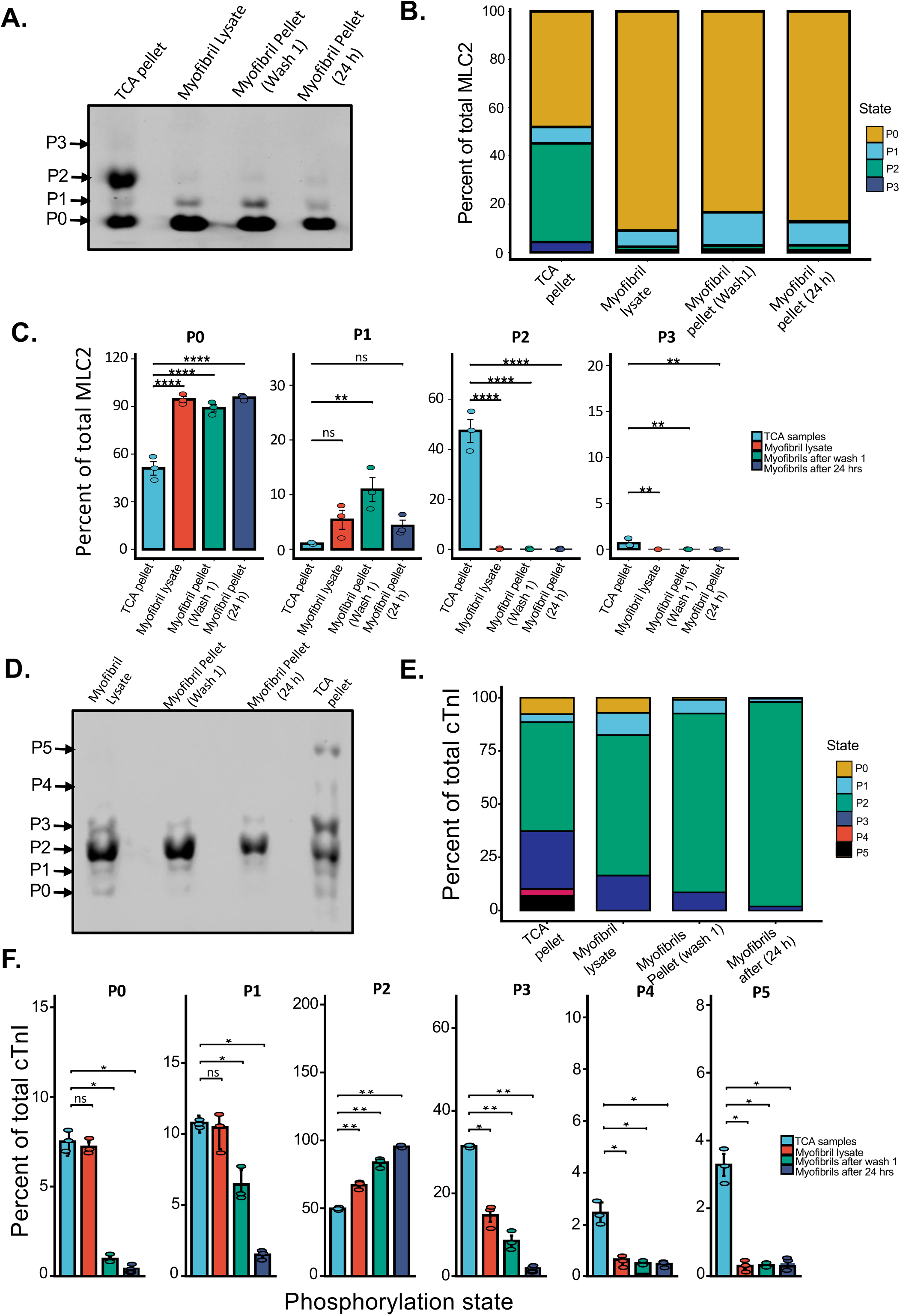
Sample preparation method alters the apparent phosphorylation-state distribution of MLC2 and cTnI in myofibrillar fractions. **(A)** Representative Mn^2+^–Phos-tag immunoblot of MLC2 phosphorylation states (P0–P3) across different sample preparation conditions: TCA-precipitated pellet, myofibril lysate, myofibril pellet (Wash 1), and myofibril pellet after 24 h. TCA-precipitated samples retained higher-order phosphorylation states, whereas myofibrillar preparations exhibited enrichment of lower phosphorylation states. **(B)** Distribution of MLC2 phosphorylation states across sample preparation conditions, expressed as the percentage of the total signal per lane. **(C)** Quantification of individual MLC2 phosphorylation states (P0–P3). TCA-precipitated samples contained significantly higher levels of the phosphorylated P2 and P3 states, whereas myofibrillar preparations showed a corresponding increase in the non-phosphorylated P0 state. Data are presented as mean ± SEM (n = 3 biological replicates); statistical comparisons are indicated (p < 0.05, p < 0.01, p < 0.001, p < 0.0001; ns, not significant). **(D)** Representative Mn^2+^–Phos-tag immunoblot of cardiac troponin I (cTnI) showing phosphorylation states (P0–P5) across the sample preparation conditions. **(E)** Distribution of cTnI phosphorylation states across sample preparation conditions, expressed as the percentage of total signal per lane. Myofibrillar preparations exhibit reduction in higher-order phosphoforms relative to TCA-precipitated samples. **(F)** Quantification of individual cTnI phosphorylation states (P0–P5) across sample preparation conditions. Myofibrillar preparations showed reduced abundance of the higher-order phosphorylation states, particularly P3–P5, together with enrichment of the P2 phosphoform. Data are presented as mean ± SEM (n = 3 biological replicates); statistical comparisons are indicated. Collectively, these findings demonstrate that sample preparation substantially influences the apparent phosphorylation-state distribution of MLC2 and cTnI, highlighting sample preparation as an important consideration for quantitative Mn^2+^–Phos-tag analysis.

## Discussion

Phos-tag SDS–PAGE has emerged as a powerful method for resolving multisite phosphorylation of proteins and has been adapted for the analysis of cardiac myofilament proteins, including MLC2, cardiac troponin I (cTnI), and cMyBP-C. The underlying Phos-tag technology was originally developed to enable phosphate-dependent mobility shifts in SDS-PAGE (11, 12, 29–31). However, substantial variability in phospho-state resolution across laboratories has limited reproducibility and hindered accurate interpretation. In the present study, we systematically optimized a Mn^2+^–Phos-tag SDS-PAGE workflow, with particular emphasis on reducing Phos-tag concentration to improve band resolution and minimize signal distortion. By evaluating gel composition, sample preparation strategies, electrophoretic conditions, and protein loading, we identified the experimental parameters that most strongly influence phospho-state separation and detection in cardiac myofilament proteins. Under these optimized conditions, Phos-tag analysis enabled robust and reproducible resolution of multiple phosphorylation states across diverse sample types. Together, these findings establish a standardized framework for high-fidelity phospho-protein analysis and highlight critical methodological variables that must be controlled for accurate interpretation of cardiac phosphorylation signaling.

### Gel chemistry dictates resolution, electrophoresis times, and blotting efficiency

Our analysis demonstrated that resolution of phosphoforms with Phos-tag gels depends on both the absolute concentration of Mn^2+^ and Phos-tag reagent as well as their concentration ratios. High Mn^2+^ concentrations (1 mM) represented a 33-fold ratio of Mn^2+^ to Phos-tag reagent, which markedly slowed migration of all proteoforms, irrespective of phosphostatus. Reducing the Mn^2+^ concentration to 300 µM (10:1 Mn^2+^: Phos-tag ratio) increased protein mobility and substantially improved phosphoform resolution. Further resolution gains were obtained by increasing the concentration of Phos-tag reagent to 50 µM (Mn^2+^/Phos-tag = 6:1). Further increasing to 75 µM (Mn^2+^/Phos-tag = 4:1) did not substantively improve resolution and, in our hands, yielded modest smearing.

The addition of Mn^2+^ and Phos-tag reagent to polyacrylamide gels, even at optimized concentrations, slows protein mobility relative to standard SDS-PAGE gels of the same acrylamide/bis-acrylamide concentrations, prompting us to compare different acrylamide concentrations. For MLC2, we found little difference in phosphoform resolution between 10% and 12% gels. We recommend 10% acrylamide, as it provides the best resolution for most myofibrillar protein targets below 50 kDa. Although we did not optimize for the resolution of cardiac myosin-binding protein C (cMyBP-C), a high-molecular-weight protein, we recommend using gels containing no more than 6% acrylamide for its analysis.

Likewise, the slow protein migration necessitates longer electrophoresis times. We evaluated constant current (25 mA/8hr; 50 mA/3hr) and constant voltage (50 V for 12 h / overnight at 4 °C). The 50 mA/3hr protocol provides an excellent option for high band separation with minimal band broadening in a reasonable amount of time. The longer overnight 50V run incurred some band broadening, however, we favored this protocol for these studies, as it appeared to maximize the detected ratio of phosphorylated to unphosphorylated protein, all else being equal. We speculate that though all electrophoresis runs were conducted at 0-4 °C localized heating under constant current may still occur, potentially impacting phosphate–Mn^2+^ coordination or contributing to further phosphate hydrolysis.

Finally, the combination of Mn^2+^ and Phos-tag reagent in the electrophoresed gel impacts electroblotting efficiency. For our studies, we opted for a traditional tank-based wet transfer protocol. The most effective transfer is achieved when electrophoresed gels are washed three times for 10 min in transfer buffer containing EDTA to chelate Mn^2+^, followed by washes in EDTA-free transfer buffer. This minimizes residual protein-Phos-tag coordination and removes Mn^2+^ ions that might impact current and heat generation upon electroblotting (12, 18).

### Attention to sample preparation minimizes Phos-tag blot artifacts while maximizing phosphoprotein detection

The unique chemistry of Phos-tag SDS-PAGE can make artifact-free protein detection more challenging when compared to standard Laemmli SDS-PAGE. Unoptimized Phos-tag protocols tend to be susceptible to band smearing and smiling, and suboptimal phosphoform resolution. This behavior can be minimized when care is taken to remove any contaminant that could impact the fidelity of Phos-tag/ Mn^2+^/Protein coordination during electrophoresis, including endogenous divalent metals and non-proteins, as well as metal chelators, including EDTA. As noted above, EDTA plays a critical role in optimal electroblotting, but its presence in the sample is detrimental to the initial electrophoresis step. Fast protein extraction from tissue into ice-cold TCA simultaneously removes contaminants that could impact electrophoresis while simultaneously arresting phosphatase and kinase activity that could impact assessments of phosphorylation. In many cases, we would also favor rapid methanol-chloroform protein extraction from tissue homogenates containing denaturants and phosphatase inhibitors. However, as discussed hereinafter, there are cases where only rapid TCA extraction maximizes phosphoform recovery. Both TCA and methanol-chloroform extraction yield insoluble protein aggregates that can be challenging to solubilize. We have found that resolubilization can be maximized by including Urea (8 M) in the sample solubilization buffer as needed. Finally, we recommend pausing electrophoresis, once samples have entered the stacking gel to flush the gel wells with electrophoresis buffer and optimize ion flow. Together, these steps substantially reduce protein migration artifacts.

It is also common practice to heat gel samples in Laemmli sample buffer at 95°C for 5 - 10 minutes to maximize protein denaturation and solubility before gel loading. However, our data indicate that standard sample-boiling protocols substantially decrease the relative abundance of protein phosphoforms. Phosphate loss is both time- and temperature-dependent. We found that heating samples at 55°C for 10 minutes mitigated phosphate loss during this step. Minimizing exposure to high temperature is also preferred if urea is present in the resolubilization buffer, to minimize protein carbamylation. It is worth noting that the importance of an optimized sample buffer heating protocol extends beyond Phos-tag applications to conventional SDS-PAGE/immunoblot protocols using phospho-specific antibodies, where maximizing the phosphoprotein signal is desirable.

### Application: assessment of myofibrillar protein phosphostatus in a myofibril preparation

With a robust Phos-tag protocol, we revisited a commonly used procedure to isolate myofibrils for activation and relaxation kinetics studies. Few myofibril preparation protocols specify the inclusion of phosphatase inhibitors that would inhibit the principal myofibril-associated phosphatases, PP1 and PP2A (22, 32–36), and, to our knowledge, none have explicitly compared the sarcomeric protein phosphorylation between TCA-homogenized hearts (the standard for preserving MLC2 phosphorylation; (37)) and isolated myofibrils. We found that in a standard phosphatase-free myofibril isolation protocol, sarcomeric proteins MLC2, cMyBP-C and cTnI all undergo dephosphorylation, albeit at different rates and to different extents. MLC2 underwent near-complete dephosphorylation, with about 90% of the protein becoming unphosphorylated by the time the heart homogenate was collected during myofibril isolation. The principal high-stoichiometry phosphorylation sites in mice are Ser 14/Ser15, and their phosphorylation is regulated by the opposing activities of cardiac myosin light chain kinase (Mylk3) and the phosphatase, PP1c (38). Ser14/15 phosphorylation is a key determinant of maximum force development (39–42), calcium sensitivity (42–45), and crossbridge cycling kinetics (40, 46). Knockout of cardiac myosin light chain kinase (Mylk3) or genetic ablation of MLC2 phosphorylation Ser14/15 in mice causes cardiac hypertrophy and impaired cardiac function (39, 41, 43, 47). Mutations in Mylk3 are associated with familial dilated cardiomyopathy (48), while downregulation loss of MLC2 phosphorylation have been reported in human (49) and experimental heart failure (50).

In contrast, we show that phosphorylation of cTnI at the high-stoichiometry sites, Ser23/24 (51, 52), regulated through the opposing actions of protein kinase A (PKA) (53–55) and PP2A (56), is relatively preserved, though higher-order lower stoichiometry phosphoforms are lost. While Ser23/24 phosphorylation may be the principal mechanism of myofibrillar desensitization and accelerated relaxation in response to b-adrenergic stimulation (55), sensitive mass spectrometry studies have identified up to 12 additional phosphorylation sites on cTnI, several of which are dysregulated in patients with ischemic heart disease or idiopathic dilated cardiomyopathy (57) and have documented impact on myofibrillar and/or cardiac function, including Tyr26 (58), Ser43/45 (59–61), Thr144 (62, 63), Ser150 (64–66), and Ser199 (67).

cMyBP-C dephosphorylation upon myofibril isolation was more extensive than that observed for cTnI, although less rapid than MLC2 dephosphorylation. The most extensively studied phosphorylation sites include the PKA-phosphorylatable Ser273, Ser282, Ser302 and Ser313 (68). This cluster constitutes an integration point for complex signaling through PKA and PKC pathways that can be offset by both PP2A and PP1C phosphatases (68, 69), impacting force generation (70), and length-dependent activation (71). The combined observations that PP2A substrate sites on cTnI are stable, and that PP1C sites on MLC2 are 90% dephosphorylated while PP1C/PP2A sites on cMyBP-C experience intermediate levels of dephosphorylation, are consistent with effective removal of PP2A activity in the early stages of myofibril preparation (i.e. multiple washes), whereas a degree of PP1C activity persists, owing to its strong interaction with thick filament-resident MYPT2(72).

The implications of protein dephosphorylation for myofibril studies will likely vary by experimental design. We have only evaluated three sarcomeric phosphoproteins and more may be affected. It can be argued that omitting phosphatase inhibitors may not grossly impact studies that address the specific impact of myofibrillar cardiomyopathy mutations, whether virally-transduced or bath-exchanged, provided that temporal batch effects are minimized. Nevertheless, tight time-matching of experiments might be required to minimize the confounding impact of progressive cMyBP-C dephosphorylation. Alternatively, inclusion of a high-affinity phosphatase inhibitor like okadaic acid in myofibril preparation buffers (e.g., (33)) would seem prudent. Tighter control and/or accounting for multi-site multi-protein dephosphorylation would be highly recommended for future studies of myofibril mechanics in the context of human heart failure and experimental heart failure models, where specific changes in MLC2, cTnI and cMyBP-C phosphorylation have been well documented.

## Conclusion

Here, we present an optimized Mn^2+^–Phos-tag SDS-PAGE analysis protocol for the analysis of select sarcomeric proteins, including a supplemental troubleshooting guide and description of the distinctions between this work and prior published protocols. We submit that the combined use of TCA-homogenized cardiac tissue and Phos-tag gels/immunoblot analysis provides a phosphorylation benchmark that can help contextualize data obtained from isolated myofibril experiments, which in turn, should help refine models of contraction that probe the relative roles of cTnI, MLC2 and cMyBP-C phosphorylation, particularly in the context of heart failure.

## Funding

This study is supported by the National Heart, Lung, and Blood Institute (NHLBI) of the National Institutes of Health (NIH) through grant R01HL164478 (DBF), and by the U.S. Army Medical Research Acquisition Activity (USAMRAA) under award HT94252410277 (DBF).

## Author Contribution

**S.B.S.** performed the experiments, analyzed the data, and wrote the manuscript. **A.F.** and **S.M.L.B.** performed myofibril preparation and reviewed the manuscript. **R.W.** contributed to cardiac tissue sample collection and reviewed the manuscript. **D.B.F.** conceived and supervised the study and contributed to data interpretation, manuscript editing, and funding acquisition.

All authors reviewed and approved the final manuscript.

---

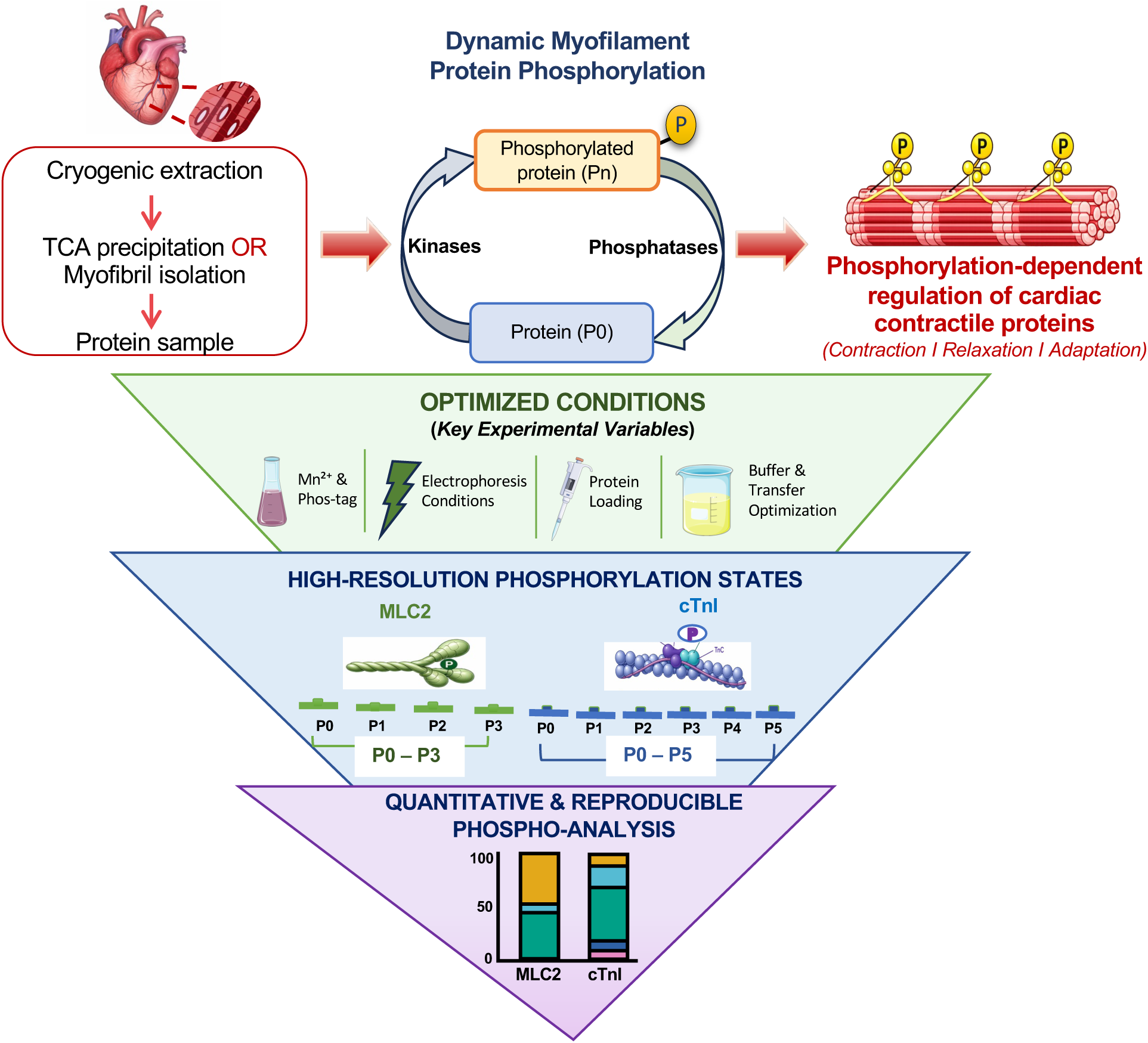

## Notes

### Competing Interest Statement

The authors have declared no competing interest.

### Summary of Updates

Table 2 has been updated to correct inadvertently repeated content in the previously uploaded version. No changes were made to the introduction, methods, results, discussion, or conclusions of the manuscript.

