## Supplementary Data for "Optimized Mn²⁺–Phos-tag Gels Reveal Sarcomeric Protein Dephosphorylation upon Myofibril Preparation"

**Supplementary Table 1**. *Key Reagents, Antibodies, and Materials Used in This Study*

| Reagent | Supplier | Catalog Number | Stock Concentration | Working Concentration | Notes |
|---|---|---|---|---|---|
| Phos-tag™ Acrylamide | FUJIFILM Wako | AAL-107 | 5 mM | 50 µM | Phosphate-affinity reagent |
| Manganese chloride ($MnCl_2$) | Sigma-Aldrich | M8266 | 30 mM | 300 µM | Metal–phosphate coordination |
| Acrylamide/Bis-acrylamide (37.5:1) | Sigma-Aldrich | A8887 / 294381 | 30% solution | 6–12% gel | Polyacrylamide gel formation |
| Tris–HCl (pH 8.8) | Thermo Fisher Scientific | BP152-1 | 1.5 M | 375 mM | Resolving gel buffer |
| Tris–HCl (pH 6.8) | Thermo Fisher Scientific | BP152-1 | 1.0 M | 125 mM | Stacking gel buffer |
| SDS (10% stock solution) | Thermo Fisher Scientific | J63394.AK | 10% | 0.1% | Protein denaturation (final concentration in gel and buffer) |
| Ammonium persulfate (APS) | Sigma-Aldrich | 248614 | 10% (fresh) | 0.05–0.1% | Polymerization initiator |
| TEMED | Sigma-Aldrich | T9281 | — | 0.05–0.1% (v/v) | Polymerization catalyst |
| Tris–glycine–SDS running buffer | Bio-Rad | 1610732 | 10× | 1× | Electrophoresis buffer (Laemmli system) |

| | | | | | |
|---|---|---|---|---|---|
| Laemmli sample buffer (2×, EDTA-free) | Bio-Rad | 1610737 | 2× | 1× | Sample preparation |
| PhosSTOP phosphatase inhibitor cocktail | Roche | 04906845001 | Tablet | 1× | Prevents dephosphorylation |
| Protease inhibitor cocktail (Complete™ Mini, EDTA-free) | Sigma-Aldrich | 11836170001 | Tablet | 1× | Prevents proteolysis |
| EDTA | Sigma-Aldrich | E7889 | 0.5 M | 30 mM | Chelates $Mn^{2+}$ to dissociate Phos-tag complexes prior to transfer |
| Glycine | Thermo Fisher Scientific | BP-381-1 | — | 192 mM | Transfer buffer component |
| Methanol | Thermo Fisher Scientific | 02-003-342 | 100% | 20% | Enhances protein binding during transfer |
| BCA protein assay | Thermo Fisher Scientific | ZA383494 | — | As per kit | Protein quantification |
| Nitrocellulose membrane | Bio-Rad | 10484059 | — | 0.2 or 0.45 µm | Protein transfer |
| Anti-MLC2 antibody | Abcam | ab92721 | — | 1:1000 | Immunoblot detection |

| | | | | | |
|---|---|---|---|---|---|
| Anti-Troponin I antibody | Sigma-Aldrich | MAB1691 | — | 1:1000 | Immunoblot detection |
| Anti-Myosin Binding Protein C (cMyBP-C / MYBPC3) | ProSci | 56-426 | — | 1:1000 | Detects MyBP-C (total or phosphorylated forms) |
| REVERT™ 700 Total Protein Stain | LI-COR | 926-11021 | — | As per manufacturer | Total protein normalization |
| Secondary anti-rabbit antibody | LI-COR | D20621-05 | — | 1:5000 | Detection |
| Secondary anti-mouse antibody | LI-COR | D31017-05 | — | 1:5000 | Detection |

### Supplementary Figure 1

**A.**

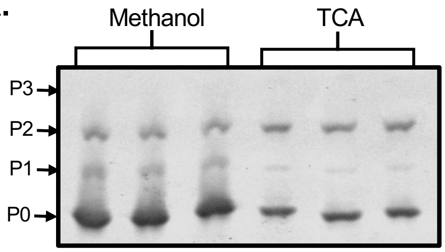


**B.**

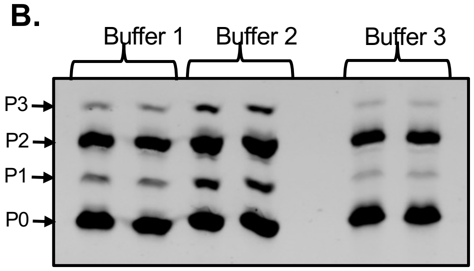


**C.** *Evaluation of three solubilization buffers for resuspending TCA pellets prior to Phos-tag SDS-PAGE.*

| Buffer | Composition | Remarks |
|---|---|---|
| **1** | 8 M urea, 100 mM NaCl, 2% SDS, 100 mM Tris-HCl (pH 8.0), 10 mM DTT | Produced complete pellet solubilization and yielded clean, well-resolved MLC2 phosphoforms. |
| **2** | 8 M urea, 100 mM Tris-HCl (pH 8.6), 23 mM glycine, 0.2 mM EDTA, 10 mM DTT | Resulted in altered migration, consistent with EDTA interference with $Mn^{2+}$–Phos-tag coordination. |
| **3** | 20% glycerol, 4% SDS, 100 mM Tris-HCl (pH 6.8), 10 mM DTT | Showed incomplete pellet solubilization and diffuse banding, indicating poor compatibility with TCA-precipitated protein pellets. |

**D.**

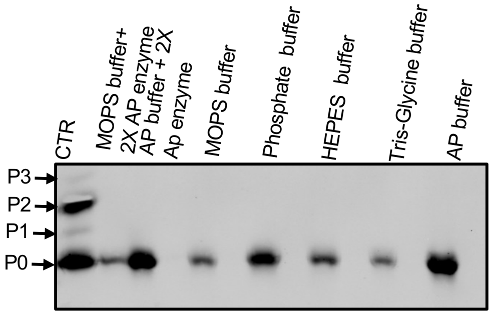


**Supplementary Figure 1. Optimization of sample preparation conditions for $Mn^{2+}$–Phos-tag analysis of MLC2. (A)** Representative $Mn^{2+}$–Phos-tag immunoblot comparing methanol/chlorofom and trichloroacetic acid (TCA) precipitation methods. TCA precipitation preserved higher-order MLC2 phosphorylation states more effectively than methanol/chlorofom precipitation. **(B)** Representative $Mn^{2+}$–Phos-tag immunoblot comparing three solubilization buffers for resuspending TCA-precipitated protein pellets before $Mn^{2+}$–Phos-tag SDS-PAGE. Buffer 1 produced complete pellet solubilization and well-resolved phosphoforms, whereas Buffer 2 resulted in altered phosphoform migration, consistent with EDTA interference with $Mn^{2+}$–Phos-tag coordination, and Buffer 3 showed incomplete pellet solubilization with diffuse banding. **(C)** Summary of the composition and performance of the three solubilization buffers evaluated for resuspending TCA-precipitated protein pellets before $Mn^{2+}$–Phos-tag SDS-PAGE. Buffer 1 provided complete pellet solubilization and optimal phosphoform resolution, Buffer 2 altered phosphoform migration, and Buffer 3 produced incomplete pellet solubilization and diffuse banding. **(D)** Representative $Mn^{2+}$–Phos-tag immunoblot evaluating the compatibility of commonly used buffer systems with MLC2 phosphorylation-state analysis, including MOPS buffer, phosphate buffer, HEPES buffer, Tris–glycine buffer, and alkaline phosphatase (AP) buffer controls. Buffer composition influenced phosphoform migration and resolution, demonstrating the importance of buffer compatibility for accurate $Mn^{2+}$–Phos-tag analysis.

#### Supplementary Figure 2

**A.**

| Loading | P0 | P1 | P2 | P3 |
|---|---|---|---|---|
| 5 µg | 74.04 | 2.57 | 21.98 | 1.41 |
| 10 µg | 70.64 | 3.00 | 24.41 | 1.94 |
| 20 µg | 68.63 | 3.24 | 26.18 | 1.95 |

**B.**

| Loading | P0 | P1 | P2 | P3 | P4 | P5 |
|---|---|---|---|---|---|---|
| 20 µg | 1.72 | 21.90 | 59.69 | 8.06 | 8.06 | 0.54 |
| 40 µg | 2.78 | 17.26 | 65.68 | 8.96 | 4.68 | 0.65 |
| 60 µg | 3.51 | 16.09 | 64.93 | 10.61 | 3.84 | 1.13 |
| 80 µg | 4.37 | 18.83 | 58.48 | 10.58 | 4.54 | 3.24 |

**Supplementary Figure 2. Quantitative summary of loading-dependent phosphorylation-state distributions for MLC2 and cTnI. (A)** Tabulated percentages of MLC2 phosphorylation states (P0–P3) at increasing protein loading amounts (5, 10, and 20 µg), expressed as the percentage of total MLC2 signal per lane. **(B)** Tabulated percentages of cTnI phosphorylation states (P0–P5) at increasing protein loading amounts (20, 40, 60, and 80 µg), expressed as the percentage of total cTnI signal per lane. These values correspond to the quantitative analyses presented in Figure 4 and illustrate the loading-dependent changes in the relative abundance of individual phosphoforms.

#### Supplementary Figure 3

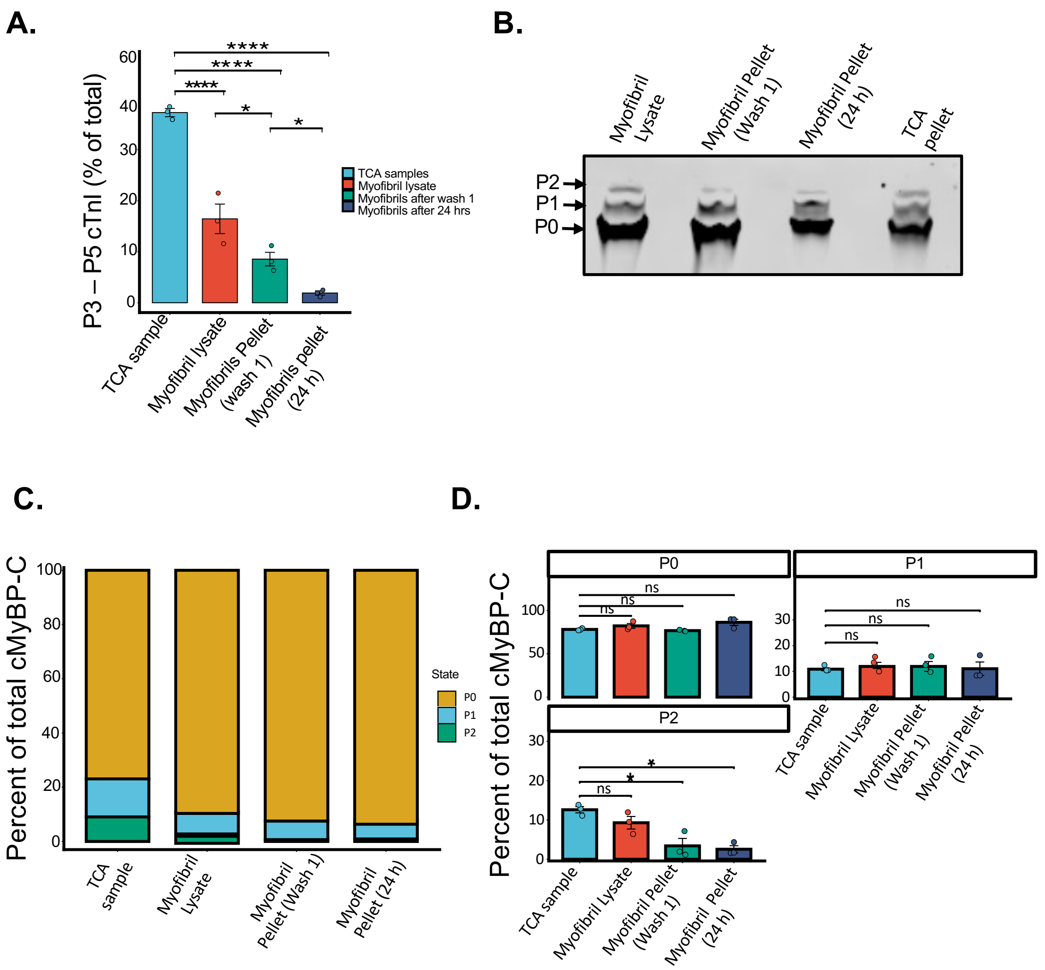


**Supplementary Figure 3. Sample preparation influences the phosphorylation-state distribution of cTnI and cMyBP-C. (A)** Quantification of higher-order cTnI phosphorylation states (P3–P5), expressed as the percentage of total cTnI signal. TCA-precipitated samples contained significantly higher levels of higher-order phosphoforms than myofibrillar preparations. Data are presented as mean ± SEM (n = 3 biological replicates), and pairwise statistical comparisons are indicated. **(B)** Representative $Mn^{2+}$–Phos-tag immunoblot of cardiac myosin-binding protein C (cMyBP-C) showing phosphorylation states (P0–P2) across the indicated sample preparation conditions. **(C)** Distribution of cMyBP-C phosphorylation states (P0–P2), expressed as the percentage of total cMyBP-C signal per lane. **(D)** Quantification of individual cMyBP-C phosphorylation states (P0–P2), presented as mean ± SEM (n = 3 biological replicates), with pairwise statistical comparisons indicated. Together, these data demonstrate that sample preparation influences the phosphorylation-state distribution of cTnI and cMyBP-C.

#### Basic steps

Optimized $Mn^{2+}$–Phos-tag SDS-PAGE workflow and critical technical considerations

$Mn^{2+}$–Phos-tag SDS-PAGE was performed as described in the main methods section, with the following optimized conditions and technical considerations identified in this study.

*1. Sample Preparation and Phosphorylation Preservation*

To preserve endogenous phosphorylation states, ventricular tissue was cryopulverized and immediately precipitated using 10% TCA containing 10 mM dithiothreitol (DTT), followed by diethyl ether washes and solubilization in urea–SDS buffer.

Importantly, avoidance of metal-chelating agents prior to electrophoresis is critical, as chelators such as EDTA or EGTA disrupt $Mn^{2+}$–Phos-tag coordination and abolish phosphorylation-dependent mobility shifts.

Myofibril isolation buffers containing EGTA were found to interfere with phospho-state resolution, resulting in altered band migration and reduced separation of higher-order phospho-species.

*2. $Mn^{2+}$–Phos-tag Gel Optimization*

Optimal separation of phospho-species was achieved using:

- Phos-tag acrylamide: 50 µM
- $MnCl_2$: 300 µM
- Acrylamide: 10% (MLC2, cTnI), 6%(cMyBP-C)

Premixing $MnCl_2$ with Phos-tag acrylamide prior to polyacrylamide gel polymerization was essential to ensure consistent metal–ligand complex formation and reproducible band resolution.

Higher $Mn^{2+}$ concentrations impaired electrophoretic mobility and compressed phospho-species, whereas lower concentrations reduced phosphate binding efficiency. Therefore, $MnCl_2$ was used in excess relative to Phos-tag to ensure efficient metal–phosphate coordination and reproducible phospho-species resolution.

*3. Electrophoresis Conditions*

Phospho-species resolution was highly sensitive to electrophoresis conditions.

Optimal separation was obtained using:

- Low-voltage, long-duration runs (50 V for 12 h/overnight at 4 °C)

Higher voltage or current conditions resulted in:

- Band broadening
- Reduced peak resolution
- Merging of closely spaced phospho-species

These effects are consistent with increased gel heating and disruption of $Mn^{2+}$–phosphate interactions.

*4. Sample Loading Effects*

Sample loading significantly influenced the apparent distribution of phospho-species.

- Higher protein loads increased signal intensity and altered the apparent abundance of individual phospho-species.
- Low-abundance phosphoforms, particularly cTnI P3–P5, were highly sensitive to protein loading, highlighting the need to optimize and standardize protein loading for accurate quantitative comparisons.

Therefore, standardized loading ranges (20–30 µg for MLC2; 40 µg for cTnI) are recommended for quantitative comparisons.

*5. $Mn^{2+}$ Removal Prior to Transfer*

Efficient transfer required complete removal of $Mn^{2+}$ ions following electrophoresis.

- Gels were incubated in 30 mM EDTA before transfer
- Followed by extensive washes in EDTA-free buffer

Incomplete $Mn^{2+}$ removal resulted in poor transfer efficiency and signal loss.

*6. Key Technical Recommendations*

Based on systematic optimization, the following conditions are critical for reproducible Phos-tag analysis:

- Use EDTA/EGTA-free buffers prior to electrophoresis
- Maintain low-temperature electrophoresis (4 °C)
- Avoid boiling samples, as excessive heat can disrupt phosphorylation-dependent mobility and reduce phospho-species resolution
- Standardize protein loading across samples
- Ensure complete $Mn^{2+}$ chelation before transfer

*7. Outcome:* Under these optimized conditions, $Mn^{2+}$–Phos-tag SDS-PAGE reliably resolved discrete phospho-species of MLC2 (P0–P3), cardiac troponin I (cTnI; P0–P5), and cardiac myosin-binding protein C (MyBPC3; three major phospho-species/bands), enabling quantitative analysis of phosphorylation patterns in cardiac tissue.
