## Supplementary material for "Optimized Mn²⁺–Phos-tag Gels Reveal Sarcomeric Protein Dephosphorylation upon Myofibril Preparation": Table 1

**Table 1:** *Comparison of Phos-tag SDS-PAGE conditions across cardiac studies*

| Study | Sample Type | Protein Target(s) | Gel (%) | Phos-tag (µM) | Mn²⁺ (µM) | Extraction / Sample Buffer | Heating Conditions | Electrophoresis Conditions | Transfer Conditions | Key Observations |
| --- | --- | --- | --- | --- | --- | --- | --- | --- | --- | --- |
| Kinoshita et al., 2009 | Purified proteins / general phosphoproteins | Multiple phosphoproteins | 8–12 | 25–100 | 25–100 | Standard SDS sample buffer | Varied | Varied | Standard wet transfer | Foundational Phos-tag methodology; establishes Mn²⁺–phosphate coordination as the basis for mobility shift |
| Messer & Marston, 2009 | Mouse/rat cardiac myofibrils | cTnI, cTnT, MLC2 | 10 | 50 | 100 | Laemmli extraction | 95°C, 5 min | 100 V constant | PVDF 0.45 µm; wet; 100 V, 1 h | First cardiac application; moderate phospho-resolution |
| Jacques et al., 2010 | Mouse ventricular tissue | cTnI | 10 | 50 | 100 | Urea + SDS buffer | 70–95°C | 100 V constant | Nitrocellulose; wet; no EDTA wash | Good P0–P2 resolution; temperature-sensitive |
| Bayliss et al., 2014 | Mouse HCM models | cTnI, cTnT | 8–10 | 40–50 | 100 | Laemmli extraction | 95°C | 120 V | PVDF (low fluorescence); wet | Lower gel % improves high-MW protein separation |
| Toepfer et al., 2013 | Mouse cardiac tissue / myofilaments | cTnI, cMyBP-C | 6–10 | 25–50 | 100 | Myofibrillar extraction | ~95°C | 100 V | Nitrocellulose or PVDF; wet transfer | Cardiac myofilament phosphorylation analysis; highlights multisite phosphorylation complexity |
| Qiu & Steinberg, 2016 | Cardiomyocyte lysates | cTnI, PLN | 10 | 40–50 | 100 | RIPA + Laemmli | 95°C | 100 V | Nitrocellulose; wet | RIPA compatible but reduces band sharpness |
| This Study (Optimized) | Mouse ventricular tissue / myofibrillar preparations | MLC2, cTnI, **cMyBP-C** | 6–12 (10% optimal for MLC2/cTnI; 6% for cMyBP-C) | 50 | 300 | TCA precipitation followed by SDS-urea sample buffer | 55°C, 10 min | 50 V, 12 h, 4 °C (MLC2); 14–16 h (cTnI). | 0.2 µm nitrocellulose; EDTA prewashes, wet; transfer 120 V, 2 h, 4°C | High-resolution phosphoform separation (P0–P5); EDTA-dependent transfer recovery; electrophoresis conditions and protein loading critically influence apparent phosphorylation stoichiometry |
