## Supplementary material for "Optimized Mn²⁺–Phos-tag Gels Reveal Sarcomeric Protein Dephosphorylation upon Myofibril Preparation": Table 2

### **Table 2: Troubleshooting Table for Phos-tag SDS–PAGE**

| Problem | Likely Cause(s) | Recommended Solution(s) |
| --- | --- | --- |
| Compressed or poorly resolved P0–Pn ladder | Excess Mn²⁺; inappropriate acrylamide percentage | Use Mn²⁺ ~300 µM and optimize gel percentage (10% for MLC2/cTnI; 6% for cMyBP-C). Note that increasing Phos-tag concentration prolongs migration time |
| Diffuse, smeared, or streaked bands | Gel overheating; sample overheating during denaturation; incomplete solubilization; excessive loading; degraded or improperly prepared buffers | Perform low-voltage electrophoresis (50 V for 12 h or ≤25 mA); ensure complete solubilization; reduce loading; use freshly prepared buffers maintained at 4°C; use freshly prepared samples. |
| Loss of phosphorylation-dependent mobility | EDTA contamination; chelating agents | Use EDTA-free reagents during sample preparation and electrophoresis. Reserve EDTA only for the post-electrophoresis wash before protein transfer. |
| Altered or unexpected phosphorylation-state distribution | Excess protein loading; phosphatase activity; sample degradation | Reduce protein loading; process samples rapidly at low temperature; include phosphatase inhibitors when appropriate; use freshly prepared samples. |
| Uneven or distorted bands | Inconsistent gel polymerization; temperature gradients; buffer variability; incorrect or inconsistent buffer pH, uneven loading; salt accumulation in wells, excessive protein concentration; incomplete protein solubilization | Standardize gel preparation and polymerization conditions; prepare and equilibrate running buffers at the correct pH; maintain consistent electrophoresis conditions; use accurate loading devices (e.g., Hamilton syringes or positive-displacement pipettes); briefly rinse or equilibrate wells with running buffer after sample loading to minimize salt and glycerol accumulation; optimize protein stock concentration to avoid overloading; and ensure complete protein solubilization before loading. |
| High background or nonspecific signal | Sample contaminants; excessive protein loading; antibody concentration too high; imaging overexposure; insufficient membrane washing | Improve sample preparation; reduce protein loading; optimize antibody concentrations and incubation times; optimize imaging settings (e.g., LI-COR); increase membrane washes (with TBST) and/or blocking time. |
| Poor resolution of higher phospho-species (P3–P5) | Excess protein loading; electrophoresis too rapid; inappropriate gel composition | Reduce protein loading; use low-voltage electrophoresis at 50 V for the optimized duration; optimize acrylamide concentration according to protein size. |
| Low signal intensity | Insufficient loading; inefficient transfer; protein degradation; low primary or secondary antibody concentration | Increase protein loading within the optimized range; optimize transfer conditions (including EDTA prewashes); perform sample preparation at low temperature; optimize primary and secondary antibody concentrations and incubation times. |
| Apparent loss of phosphorylation | Phosphatase activity; delayed sample processing; heat generation during sample preparation or electrophoresis. | Process samples rapidly; maintain samples on ice; include phosphatase inhibitors where appropriate; avoid excessive heating during denaturation and electrophoresis; use the optimized denaturation condition (55°C for 10 min). |
| High inter-experimental variability | Reagent variability; EDTA contamination; inconsistent electrophoresis conditions; incomplete sample solubilization. | Use consistent reagent batches; standardize all experimental conditions; verify buffer pH; ensure complete sample solubilization before loading; maintain identical electrophoresis and transfer conditions across experiments. |
| Uneven electrophoretic migration across lanes | Unstable or malfunctioning power supply; poor electrode contact; damaged electrophoresis apparatus; uneven buffer levels | Verify stable voltage/current output before each run; inspect power cables and electrodes for secure connections; ensure equal buffer levels in both chambers; replace or service faulty electrophoresis equipment if necessary. |
| **Saturated phospho-species signal** | **High-abundance phospho-species exceeds the linear detection range** | **Reimage the blot at a lower acquisition intensity before quantitative analysis.** Ensure all bands used for densitometry are within the linear detection range. |
